# Arachidonic acid modulates fluconazole-induced responses in *Candida albicans* biofilms

**DOI:** 10.1101/2025.11.25.690624

**Authors:** Ruan Fourie, Oluwasegun Kuloyo, Chibuike Ibe, Gabre Kemp, Previn Naicker, Ireshyn Govender, Michael Kruger, Olihile Sebolai, Jacobus Albertyn, Carolina Pohl

## Abstract

*Candida albicans* is a human commensal, which causes opportunistic infections with high morbidity and mortality and is characterised by the production of resistant biofilms. Polyunsaturated fatty acids, such as arachidonic acid, can increase azole susceptibility of *C*. *albicans* biofilms significantly. However, the underlining mechanism is not known. We investigated the effect of arachidonic acid on known fluconazole resistance mechanisms namely, overexpression of *ERG11*, increased ergosterol content, oxidative stress resistance, as well as expression and activity of the efflux pump, Cdr1p. Upregulation of *ERG11* was observed in biofilms exposed to fluconazole. However, this was reversed in the presence of arachidonic acid, even in the presence of fluconazole. Furthermore, arachidonic acid downregulated the fluconazole-induced oxidative stress response of *C. albicans*. Previous transcriptome data indicated a significant increase in *CDR1* expression during early biofilm formation in the presence of arachidonic acid. However, we found that efflux activity was reduced in the presence of arachidonic acid, which indicates the loss of function of the membrane-associated protein. This contradictory phenomenon was further investigated by determining the effect of arachidonic acid on the localisation and phosphorylation of Cdr1p (ProteomeXchange identifier PXD070958). Our results show that the presence of arachidonic acid may cause mislocalisation and changes in the phosphorylation of Cdr1p; all of which could impact its activity. These results demonstrate that multiple mechanisms are potentially involved in the arachidonic acid-induced increased fluconazole susceptibility and allows for further exploration of lipid-mediated modulation of antifungal responses.

## INTRODUCTION

*Candida albicans* is the primary cause of the life-threatening systemic candidiasis in immunocompromised individuals, transplant recipients and patients undergoing chemotherapy (Shareck and Belhumeur, 2011). An essential virulence factor of *C. albicans* is the formation of biofilms on abiotic and biotic surfaces (Mayer et al., 2013; Sardi et al., 2013). These biofilms are notorious for their high antifungal resistance, and serve as a potential source of systemic infection (Gulati and Nobile, 2016; Tsui et al., 2016). Although several antifungal classes have been identified as treatment options for *C. albicans* infections, the azoles, particularly fluconazole, is the most administered, due to excellent bioavailability, low cost and reduced toxicity (Lockhart and Berkow, 2017). Fluconazole acts by inhibiting the lanosterol 14-α demethylase (encoded by *ERG11*) which catalyses the oxidative C14-demethylation of lanosterol during the synthesis of fungal-specific ergosterol (Shapiro et al., 2011; Tobudic et al., 2012). Fluconazole also induces oxidative stress in *C. albicans* by the generation of reactive oxygen species (ROS) (Kobayashi et al., 2002; Wang et al., 2009; Arana et al., 2010), which may damage proteins, lipids (via lipid peroxidation) as well as nucleic acids, and stimulates apoptosis (Farrugia and Balzan, 2012). However, high resistance, especially in the treatment of biofilm-associated infections, hampers its efficiency (Taff et al., 2013; Fothergill et al., 2014; Whaley et al., 2017).

Several resistance mechanisms, including the increased expression of *ERG11*, alteration of sterol synthesis and overexpression of drug efflux pumps are employed by *C. albicans* against fluconazole (Berkow and Lockhart, 2017; Whaley et al., 2017). In addition, C. albicans has . albicans has robust antioxidant defence mechanisms to overcome oxidative attack and which have been reported to be upregulated in fluconazole resistant isolates (Poopedi et al., 2021).

Arachidonic acid (*cis*-5,8,11,14-eicosatetraenoic acid) is an ω-6 C20 fatty acid present in mammalian cells, esterified to membrane phospholipids in the sn-2 position (Hanna and Hafez, 2018), from where it can be released through the action of phospholipase A2 (PLA2) (Rodríguez *et al*., 2014). The release of arachidonic acid can be stimulated by bacterial lipopolysaccharide and fungal cell wall components, such as β-(1,3)-glucans. In addition, *C. albicans* can directly release arachidonic acid from host cell membranes by the action of its own PLA2 enzyme (Castro et al., 1994). Although the oxygenated products of arachidonic acid are well known to be involved in the inflammatory response towards infection (Dennis and Norris, 2015), the non-oxygenated fatty acid may also play important roles during infection and is proposed to be an endogenous antimicrobial molecule, released by immune cells as a defence mechanism against invading pathogens (Das, 2018). Arachidonic acid has been shown to inhibit morphogenesis of *C. albicans* (Shareck and Belhumeur, 2011; Kuloyo et al., 2020) and to increase the susceptibility of *C. albicans* biofilms to fluconazole (Ells et al., 2009). However, the mechanism(s) by which arachidonic acid influences fluconazole susceptibility is unclear. Therefore, the aim of this study was to examine the influence of arachidonic acid on different fluconazole responses of *C. albicans* biofilms.

## MATERIALS AND METHODS

### Strains used

The strains used in this study are summarized in Table 1 and were stored at -80°C with 15% glycerol. Yeast strains were revived and maintained on Yeast-Malt extract (YM) agar (3 g/L malt extract, 3 g/L yeast extract, 5 g/L peptone, 10 g/L glucose and 16 g/L agar) at 30°C.

**Table 1.** Strains used in this study.

| Strain | Genotype | Parental strain | Reference |
| --- | --- | --- | --- |
| SC5314 | Wild type | N/A | N/A |
| CAI4 | *ura3::imm434/ura3::imm434 iro1/iro1::imm434* | CAF2-1 | Fonzi and Irwin (1993) |
| ASCa1 | *ura3Δ::imm434/URA3 CDR1-GFP* | CAF4-2 | Szczepaniak *et al.* (2015) |
| *cdr1Δ/Δ* | *cdr1Δ;cdr1Δ* | SC5314 | This study |
| *cdr1Δ/Δ::CDR1* | *cdr1Δ::CDR1;cdr1Δ::CDR1* | SC5314 | This study |
| *gpx2Δ* | *tetO-GPX2/gpx2Δ* | CaSS1 via CAI4 | Roemer *et al.* (2003) |
| *prx1Δ* | *tetO-PRX1/prx1Δ* | CaSS1 via CAI4 | Roemer *et al.* (2003) |
| *pst1Δ* | *tetO-PST1/pst1Δ* | CaSS1 via CAI4 | Roemer *et al.* (2003) |
| *sod5Δ* | *tetO-SOD5/sod5Δ* | CaSS1 via CAI4 | Roemer *et al.* (2003) |
| *tps1Δ* | *tetO-TPS1/tps1Δ* | CaSS1 via CAI4 | Roemer *et al.* (2003) |
| *zwf1Δ* | *tetO-ZWF1/zwf1Δ* | CaSS1 via CAI4 | Roemer *et al.* (2003) |

### Biofilm formation

All *C. albicans* strains were grown on YM extract agar for 24 h at 30°C and were inoculated into 10 mL yeast nitrogen base (YNB) broth (10 g/L glucose and 16 g/L YNB) and incubated at 30°C for 24 h. Cells were harvested at 1878 *g* for 5 min and the supernatant removed. This was followed by washing the cells twice with phosphate buffered saline (PBS) (Oxoid, England). The cells were then counted with a haemocytometer and diluted to 1 × 10^6^ cells/mL in 20 mL filter sterilized (0.22 µm nitrocellulose filter, Merck Millipore, Ireland) RPMI-1640 medium (Sigma-Aldrich, USA) dispensed into either 90 mm polystyrene petri dishes, 6-well plates, or 96-well plates (Merck, Germany) and incubated for either 6 h (for immature biofilms) or 48 h (for mature biofilms) at 37°C as indicated, to allow biofilm formation.

### Expression levels of *ERG* genes and ergosterol quantification

Transcriptome data from Kuloyo and co-workers (2020) (NCBI GEO accession number GSE137423), obtained for *C albicans* SC5314 6-h biofilms, were investigated for gene expression levels of *ERG* genes in the presence of either 1 mM arachidonic acid, 1 mg/L fluconazole, or a combination of the two compounds compared to solvent controls, containing less than 1% solvent.

Ergosterol was extracted from *C*. *albicans* SC5314 biofilms as described (Arthington-Skaggs *et al*., 1999), with slight modifications. Immature (6 h) biofilms were prepared in the presence of either 1 mM arachidonic acid, 1mg/L fluconazole or a combination of the two compounds. The biofilms were harvested by centrifugation at 2700 *g* for 5 min and washed with sterile distilled water. The biofilms were placed in a borosilicate tube while 3 mL of freshly prepared potassium hydroxide (25% m/v), dissolved with a solution of methanol, ethanol and water (700:315:15), was added. The solution was heated to 85°C for 1 h in a water bath and allowed to cool to room temperature. Ergosterol was extracted with the addition of 3 mL of n-heptane, containing 10 μg/mL of cholesterol internal standard and 1 mL of water. The solution was mixed with a vortex and allowed to separate without disruption for 30 min. The n-heptane layer was collected into a new borosilicate tube and stored in -20°C. Derivatisation was done by evaporating the n-heptane to dryness under a stream of nitrogen gas and the addition of 50 μL of N,O-bis(trimethylsilyl)acetamide containing 2% trimethylchlorosilane and incubation for 18 h at 80°C. After incubation, the solution was evaporated to dryness under a stream of nitrogen gas and resuspended in 100 μL of n-decane. The GC/MS analysis was carried out on a Thermo Electron Trace 1310 Gas chromatograph connected to an ISQ 7000 single quadrupole mass spectrometer. Separation was achieved with an Agilent FactorFour VF-5 column with the following dimensions: length of 60 m, inner diameter of 0.32 mm, and a film thickness of 0.25 μm.

Helium was used as the carrier gas at a flow rate of 5 mL/min. The injection port temperature was 175°C with a split ratio of 33:1. The initial oven temperature was held at 175°C for 1 min. The temperature was subsequently raised at a rate of 7°C/min until 300°C was achieved, and held for 10 min. Electron impact ionization was employed. The MS source temperature was 250°C. Mass analysis was carried out in the specific ion monitoring mode. Data was captured and analysed with Thermo Xcalibur 4.0 software employing the NIST 17 spectral library.

### Determination of the influence of arachidonic acid on oxidative stress

Transcriptome data from Kuloyo and co-workers (2020) obtained for *C albicans* SC5314 6-h biofilms, were investigated for differentially regulated genes involved in response to oxidative stress. *Candida albicans* heterozygous mutants corresponding to selected genes, exhibiting changes in the levels of differential regulation in the presence of fluconazole and arachidonic acid, were selected from a double-barcoded library of heterozygous *C. albicans* mutants (version 2013A) supplied by Merck Sharp & Dohme Corp (Table 1) (Roemer *et al*., 2003). Mature biofilms were grown from the mutants, *C. albicans* CA14 as well as *C. albicans* SC5314 for 48 h in the presence of 1 mg/L fluconazole at 37°C for 48 h. Biofilm metabolic activity was measured by assessing formazan formation through the reduction of 2,3-bis (2-methoxy-4-nitro-5-sulfophenyl)-5-[(phenylamino) carbonyl]-2H-tetrazolium hydroxide (XTT) by mitochondrial dehydrogenases as described by Kuhn and co-workers (2003). In addition, 0.045 mM butylated hydroxytoluene (BHT) was added to the biofilms of a selected mutant (*pst1Δ*) and inhibition by fluconazole determined.

Biofilms of *C. albicans* SC5314, *cdr1*Δ/Δ, as well as the add-back strain (*cdr1*Δ/Δ ::*CDR1*) were also prepared in 96-well plates as described above in the absence or presence of 0.1 mM arachidonic acid. In addition, 0.5 mM α-tocopherol or 0.045 mM BHT were added, and biofilm biomass was quantified with the crystal violet assay (Jin et al., 2003). Briefly, the supernatant from each well was removed and the biofilms were washed twice with sterile PBS. Biofilms were then left to air dry at room temperature for 45 minutes and stained with 110 μL crystal violet (0.4 % w/v) for 45 min. Biofilms were washed three times with 350 μL sterile H_2_O and de-stained with 200 μL 95 % ethanol for 45 min. One hundred μL of de-staining solution was transferred to a clean 96-well plate and absorbance measured at 595 nm.

### Expression levels of efflux pump genes and activity assay

Transcriptome data from Kuloyo and co-workers (2020), obtained for *C albicans* SC5314 immature (6 h) biofilms, were investigated for differential expression of genes encoding for drug efflux pumps in the presence of either 1 mM arachidonic acid, 1mg/L fluconazole, or a combination of the two compounds, compared to solvent controls.

Uptake and glucose-induced efflux of rhodamine 6G (Rh6G) by *C*. *albicans* SC5314 biofilms were conducted as previously described (Maesaki et al., 1999). Briefly, immature (6 h) biofilms were prepared in black flat-bottomed 96-well microtiter plates. Afterwards, the media was removed, and the biofilms were washed twice with sterile PBS. The biofilms were incubated in sterile PBS to de-energise for 1 h at 37°C. The biofilms were incubated in 10 μM Rh6G in dimethyl sulfoxide (DMSO) at 37°C for 1 h. Fluorescence was measured at an excitation of 530 nm and an emission of 590 nm, every 10 min for 60 min, to evaluate the uptake of Rh6G. After Rh6G uptake measurements, the biofilms were washed with sterile PBS and treated with either 0.1 mM or 1 mM arachidonic acid. An additional experiment was conducted where arachidonic acid was added at the start of biofilm formation. Rh6G efflux from the cells was induced with the addition of 2 mM glucose in PBS. Fluorescence measurement was repeated to evaluate the efflux of Rh6G.

### Fluorescence microscopy

*Candida albican*s ASCa1, containing a CDR1-GFP fusion (Szczepaniak *et al*., 2015) was used to prepare immature (6 h) biofilms in 6-well culture plates in the presence of arachidonic acid (1 mM) or equivalent ethanol control. After biofilm formation, cells were scraped off and re-suspended in paraformaldehyde (4 g/L), incubated at room temperature for 15 min, followed by washing of cells in with a potassium phosphate/sorbitol mixture (1.2 M sorbitol with 0.1 M potassium phosphate). Excitation intensity was kept identical for the different samples at the peak excitation wavelength corresponding to fluorescence of GFP (488 nm) and resultant fluorescence was visualised (Nikon TE 2000, Japan).

### Determination of phosphorylation of Cdr1p

Cells of *C. albicans* SC5314 were grown as planktonic cultures as well as biofilms. A single yeast colony was grown in YPD overnight, diluted to an OD_600_ of 0.1 and further allowed to grow to mid-logarithmic phase. The cultures were then exposed to 1 mM arachidonic acid, 1mg/L fluconazole, or solvent control for 1 h. After treatment, the cells were washed twice in ice-cold sterile PBS and pelleted at 4200 *g* for 5 min. In addition, biofilms were prepared as described for 6 h in the presence of 1 mM arachidonic acid, 1mg/L fluconazole, or solvent control, harvested, and washed twice with PBS.

For both types of cultures, plasma membranes were extracted according to Madani and co-workers (2021) with slight modification. Briefly, the cell pellets from planktonic cultures as well as biofilms were resuspended in 0.5 mL homogenizing buffer (HB) (50 mM Tris, 0.5 mM EDTA, 20% (v/v) glycerol; pH 7.5) containing PhosSTOP phosphatase inhibitor cocktail tablet (Roche) and 1 mM phenylmethylsulfonyl fluoride (PMSF), and snap frozen in liquid nitrogen. The frozen cells were broken using sterile mortar and pestle, collected and centrifuged at 5200 *g* for 5 min at 4°C to remove cell debris, unbroken cells, and nuclei. The supernatant (450 µL) was collected in a microcentrifuge tube containing 1 mL of ice-cold HB supplemented with fresh PMSF (1 mM) and centrifuged at 18000 *g* for 1 h at 4°C. The pellets were resuspended in 100 µL HB and the concentration of protein determined using the Pierce BCA protein assay as per manufacturer’s instruction.

Protein samples (35 μg per sample) were reduced with 5 mM tris(2-carboxyethyl)phosphine and alkylated with 10 mM 2-chloroacetamide at room temperature for 20 min. Thereafter, samples were purified of detergents and salts by protein aggregation capture using MagReSyn™ Hydroxyl beads (ReSyn Biosciences) as previously described (Batth *et al*., 2019; Baichan *et al*. 2023). On-bead protein digestion was performed using a 1:40, protease:protein, ratio of sequencing-grade trypsin for 4 h at 47°C. Peptides were desalted using a Sep-Pak C18 40 mg 96-well plate and eluted in 500 μL of 60% acetonitrile. Eluates were dried to near completion (<50 μL) using a vacuum concentrator. Thereafter, eluates were immediately diluted in 500 μL of phosphopeptide enrichment binding buffer (1M glycolic acid in 80% acetonitrile and 5% trifluoroacetic acid). Phosphopeptide enrichment was performed using MagReSyn™ Zr-IMAC HP beads (ReSyn Biosciences) as previously described (Tape et al., 2014; Arribas Diez *et al*., 2021). Eluted phosphopeptides were dried and stored at -80°C until LC-MS analysis.

Phosphopeptides were resuspended in 0.1% formic acid (Solvent A), loaded onto Evotips and analyzed using an Evosep One LC system coupled to a Bruker sims-TOF HT mass spectrometer. Injected peptides were gradient-eluted and separated on a PrepSep C18 column (75 μm × 15 cm, 1.9 µm particle size) with a preformed acetonitrile gradient over 30 min (40 SPD method). The Bruker sims TOF HT mass spectrometer was operated in DIA-PASEF mode. Precursor peptides were selected in the 475-1150 m/z range and 0.817-1.315 1/K0 ion mobility range. This was done by using 27 fixed m/z windows of 25 Da each with no overlap between windows. Fragment ions were acquired from 100-1700 m/z during the total cycle time of 1.065 s.

DIA data was processed using Spectronaut v19 software (Biognosys). The default direct DIA identification and quantification settings were used for data processing. Carbamidomethylation was added as a fixed modification. Phosphorylation of serine, threonine, and tyrosine, N-terminal acetylation, and methionine oxidation were added as variable modifications. The reference proteome for *C. albicans* strain SC5314 (downloaded on 27 June 2024 from www.uniprot.org) and common contaminating proteins was used as the search database. A q-value ≤ 0.01 cut-off was applied at the precursor and protein levels. Quantification was performed at the MS1 and MS2 level. Label-free cross-run normalisation was employed using a global normalisation strategy. Peptides were collapsed according to their phosphorylation site in Spectronaut to generate a PTM site report. The mass spectrometry proteomics data have been deposited to the ProteomeXchange Consortium via the PRIDE (Deutsch et al., 2023; Perez-Riverol et al., 2025) partner repository with the dataset identifier PXD070958. Candidate dysregulated phosphorylation sites were filtered at a q-value ≤ 0.05 and absolute Log_2_ fold change (FC) ≥ 0.58.

### Statistical analyses

All quantitative experiments were conducted in biological triplicates, each with at least three technical replicates, and the averages and standard deviations were calculated. The t-test was performed in order to assess the significance of differences. *P* values of 0.05 or less were considered to be significant.

## Results

### Arachidonic acid overrides the upregulation of *ERG11* by fluconazole

According to Kuloyo and co-workers (2020), fluconazole was seen to cause a modest upregulation in the expression of several genes involved in ergosterol synthesis (Table 2). Most of these genes were not differentially regulated by arachidonic acid alone, with the notable exception of *ERG11*, which was slightly downregulated. Interestingly, when arachidonic acid was combined with fluconazole, the expression of *ERG1*, *ERG4,* and *ERG11* were returned to control levels. Figure 1 indicates the ergosterol levels of cells exposed to fluconazole and arachidonic acid, relative to the control cells. As expected, and despite the upregulation of *ERG* genes by fluconazole, the level of ergosterol in the cells exposed to fluconazole was lower than in the control (Mukherjee *et al*., 2003; Ramage *et al*., 2012; Chandra and Mukherjee, 2015). Similar to work by Ells and co-workers (2009), an increase in ergosterol content of *C. albicans* biofilms treated with arachidonic acid was observed, however the combination of fluconazole and arachidonic acid also caused a decrease in ergosterol. This suggests that the presence of arachidonic acid may override the fluconazole-induced upregulation of *ERG11*, but does not impact the fluconazole-induced decrease in ergosterol levels.

**Figure 1.**
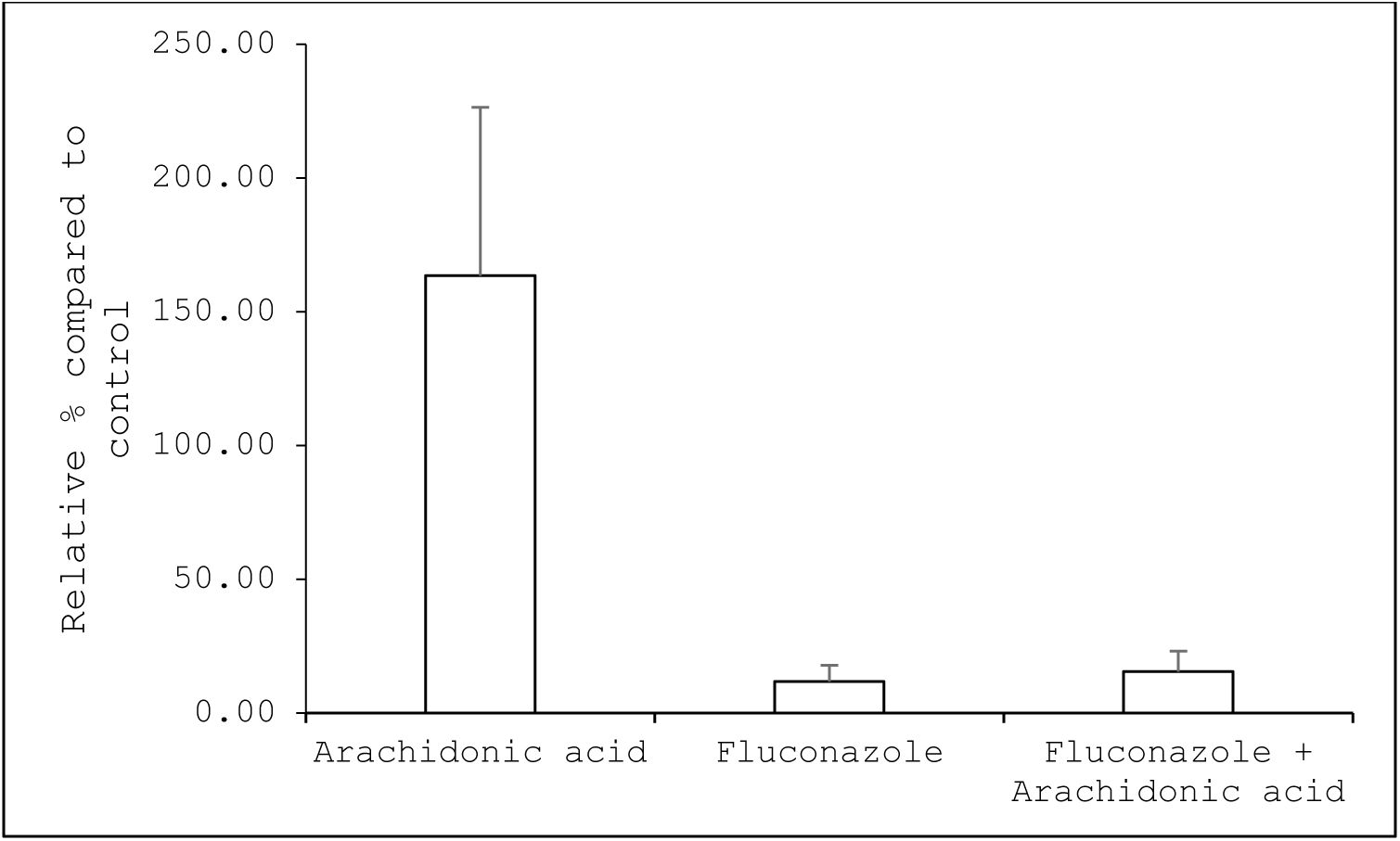
Effect of fluconazole and arachidonic acid on the ergosterol levels of *C. albicans* biofilm. Values are the mean of three independent experiments, and the standard deviation is indicated by the error bars.

**Table 2.** Differential expression of genes involved in ergosterol synthesis.

| Gene | Log<sub>2</sub>fold change in the presence of different compounds |  |  |
| --- | --- | --- | --- |
|  | Fluconazole | Arachidonic acid | Fluconazole + arachidonic acid |
| *ERG1* | 1.7 | -0.7 | NDE* |
| *ERG2* | NDE | NDE | 1.1 |
| *ERG3* | 1.0 | NDE | -0.6 |
| *ERG4* | 0.8 | NDE | NDE |
| *ERG5* | 0.9 | NDE | 0.8 |
| *ERG6* | 2.8 | NDE | 2.3 |
| *ERG7* | 0.7 | NDE | 0.7 |
| *ERG9* | 1.3 | NDE | 0.6 |
| *ERG11* | 1.3 | -1.1 | NDE |

\* NDE – Not differentially expressed compared to solvent control

### Arachidonic acid influences fluconazole-induced oxidative stress responses

Fluconazole is known to induce the formation of ROS in *C. albicans* (Arana *et al*. 2010). However, in response to ROS, *C. albicans* expresses oxidative stress response and antioxidant systems such as *CAT1*, *TRX1*, *GRX2*, *SOD1* and *SOD5* (Abegg *et al*., 2011; Dantas *et al*., 2015; Komalapriya *et al*., 2015; Poopedi et al. 2021). Data from Kuloyo and co-workers (2020) showed the differential expression of genes involved in the response to oxidative stress (Table 3).

**Table 3.** Differential expression of genes associated with response to oxidative stress in *Candida albicans* exposed to fluconazole, arachidonic acid, or a combination of the two compounds.

| Genes | Description | Log<sub>2</sub>fold change |  |  | Δ* |
| --- | --- | --- | --- | --- | --- |
|  |  | Fluconazole | Arachidonic acid | Fluconazole + Arachidonic acid |  |
| *GPX2* | Putative glutathione peroxidase | 0.3 | NDE** | -2.6 | -2.9 |
| *SOD5* | Cu-containing superoxide dismutase | 0.8 | NDE | -1 | -1.8 |
| *PST1* | Flavodoxin-like protein | -0.8 | -1.3 | -2.4 | -1.6 |
| *PRX1* | Thioredoxin peroxidase | NDE | NDE | -1.1 | -1.1 |
| *TPS1* | Trehalose-6-phosphate synthase | NDE | -0.6 | -1.1 | -1.1 |
| *ZWF1* | Glucose-6-phosphate dehydrogenase | NDE | -0.7 | -1.1 | -1.1 |
| *SUR7* | Required for normal cell wall, plasma membrane, cytoskeletal organization, endocytosis | 0.6 | NDE | -0.5 | -1.1 |
| *SOH1* | Subunit of the RNA polymerase II mediator complex | NDE | -0.9 | -1 | -1 |
| *HSP21* | Small heat shock protein | 0.9 | 0.9 | NDE | -0.9 |
| *GPX1* | Putative glutathione peroxidase | NDE | NDE | -0.7 | -0.7 |
| *MDR1* | Plasma membrane MDR/MFS multidrug efflux pump | 1.2 | NDE | 0.5 | -0.7 |
| *orf19.2782* | Ortholog(s) have 2 iron, 2 Sulphur cluster binding, disulfide oxidoreductase activity, glutathione disulfide oxidoreductase activity | 0.7 | NDE | NDE | -0.7 |
| *DDR48* | Immunogenic stress-associated protein | 2.5 | 1.1 | 1.9 | -0.6 |
| *PST2* | Flavodoxin-like protein | -0.7 | NDE | -1.3 | -0.6 |
| *GLX3* | Glutathione-independent glyoxalase | -0.6 | -0.8 | -1.2 | -0.6 |
| *AFT2* | Putative Aft domain transcription factor | -0.4 | NDE | -1 | -0.6 |
| *HSP12* | Heat-shock protein | 0.6 | 1 | NDE | -0.6 |
| *CHK1* | Histidine kinase | 1.1 | 0.5 | 0.6 | -0.5 |
| *SSU81* | Predicted adaptor protein involved in activation of MAP kinase-dependent signalling pathways | NDE | NDE | -0.5 | -0.5 |
| *RSV167* | SH3-domain- and BAR domain-containing protein involved in endocytosis | 0.5 | NDE | NDE | -0.5 |
| *CPX31* | Putative glutathione peroxidase | 0.5 | -1.3 | NDE | -0.5 |
| *ARF1* | Golgi-localized ADP-ribosylation factor | -0.7 | -0.6 | -1.2 | -0.5 |
| *TPS2* | Trehalose-6-phosphate phosphatase | -0.5 | NDE | -1 | -0.5 |
| *HVN1* | Modulates virulence and antioxidant defence pathways | 2.6 | 1.4 | 2.2 | -0.4 |
| *RAS1* | RAS signal transduction GTPase | -0.7 | NDE | -1 | -0.3 |
| *TTR1* | Putative glutaredoxin | 1.6 | NDE | 1.4 | -0.2 |
| *YCP4* | Flavodoxin-like protein | 1.6 | 1.3 | 1.5 | -0.1 |
| *DSE1* | Essential cell wall protein | 0.4 | 0.8 | 0.3 | -0.1 |
| *MED3* | Subunit of the RNA polymerase II mediator complex | -0.6 | -0.6 | -0.7 | -0.1 |
| *orf19.5276* | Putative nuclear pore-associated protein | 0.7 | NDE | 0.6 | -0.1 |
| *SAC6* | Fimbrin | 1.7 | 0.6 | 1.6 | -0.1 |
| *HST3* | Histone H3K56 deacetylase | -1 | -0.6 | -1 | 0 |
| *HOG1* | MAP kinase of osmotic-, heavy metal-, and core stress response | -0.6 | NDE | -0.6 | 0 |
| *PBS2* | MAPK kinase | -0.6 | -0.3 | -0.6 | 0 |
| *TPK1* | cAMP-dependent protein kinase catalytic subunit | 1.9 | 1.6 | 1.9 | 0 |
| *DBP4* | Putative DNA polymerase epsilon subunit D | -1 | -0.9 | -1 | 0 |
| *CKB1* | Regulatory subunit of protein kinase CK2 | -0.6 | NDE | -0.5 | 0.1 |
| *FDH3* | Glutathione-dependent formaldehyde dehydrogenase | 0.7 | NDE | 0.8 | 0.1 |
| *SRR1* | Two-component system response regulator | 1.5 | 0.9 | 1.7 | 0.2 |
| *RAD52* | Required for homologous DNA recombination | -0.8 | -0.6 | -0.6 | 0.2 |
| *SLN1* | Histidine kinase | -0.9 | NDE | -0.6 | 0.3 |
| *SOD4* | Cu-containing superoxide dismutase | 1 | NDE | 1.3 | 0.3 |
| *TAC1* | Zn(2)-Cys(6) transcriptional activator | 1.5 | 1.1 | 1.8 | 0.3 |
| *SFP1* | C2H2 transcription factor | 0.6 | 0.6 | 0.9 | 0.3 |
| *MMS21* | Putative SUMO E3 ligase | -0.4 | NDE | NDE | 0.4 |
| *MPS1* | Monopolar spindle protein | -0.9 | NDE | -0.5 | 0.4 |
| *PDE2* | High affinity cyclic nucleotide phosphodiesterase | 0.9 | 1.3 | 1.4 | 0.5 |
| *NRG1* | Transcription factor/repressor | 1.6 | 1.9 | 2.1 | 0.5 |
| *PST3* | Flavodoxin-like protein | 2.2 | 1.9 | 2.8 | 0.6 |
| *STH1* | Putative ATP-dependent helicase | -0.6 | 0.3 | NDE | 0.6 |
| *TRX1* | Thioredoxin | 1.2 | 1.6 | 1.9 | 0.7 |
| *HOF1* | Involved in cytokinesis and DNA damage response | -2.5 | NDE | -1.6 | 0.9 |
| *ALO1* | D-Arabinono-1,4-lactone oxidase | -0.9 | NDE | NDE | 0.9 |
| *RAP1* | Transcription factor | -0.9 | NDE | NDE | 0.9 |
| *OGG1* | Mitochondrial glycosylase/lyase | -1 | NDE | NDE | 1 |
| *YDJ1* | Putative type I HSP40 co-chaperone | 1 | 0.5 | 2.4 | 1.4 |
| *RAD51* | Involved in homologous recombination and DNA repair | -1.7 | NDE | NDE | 1.7 |
| orf19.2175 | Mitochondrial apoptosis-inducing factor | 0.8 | 3.1 | 3.1 | 2.3 |

\*Change in differential expression in the presence of both fluconazole and arachidonic acid versus fluconazole alone
\*\*NDE – Not differentially expressed compared to solvent control

Although some of the genes associated with response to oxidative stress did not show changes in differential expression when arachidonic acid was added to fluconazole, orf19.2175 (C2_08100W_A), which encodes a mitochondrial apoptosis inducing factor (Ma *et al*., 2016), was upregulated in the presence of arachidonic acid, compared to fluconazole alone. In addition, *OGG1*, which plays a role in repairing oxidative damage to mitochondrial DNA (Li et al., 2017), *YDJ1*, which plays a role in mitochondrial protein import (Xie et al., 2017) and *RAD51* which plays a role in DNA repair (Kumari et al.,2023) are also upregulated. These increased expression levels suggest the ability of arachidonic acid to cause apoptosis in *C. albicans*, similarly to other polyunsaturated fatty acids (Thibane *et al*., 2012).

In addition, many genes were significantly downregulated by the combination compared to fluconazole alone, suggesting that arachidonic acid influences the ability of *C. albicans* to respond to fluconazole-induced oxidative stress. To confirm that these genes are important in the response to fluconazole, the susceptibility of heterozygous mutants, containing only one copy of the selected gene, compared to their parental strain and *C. albicans* SC5314 was investigated. All the mutants displayed a significant reduction in metabolic activity in the presence of fluconazole, compared to the wild type and parental strain (Figure 2a). The heterozygous *pst1Δ* mutant was selected as a representative to test for the reversal of fluconazole-mediated oxidative stress in the presence of the antioxidant, BHT. The addition of BHT partially reversed the effect of fluconazole on *pst1Δ* (Figure 2b). A similar reversal of fluconazole susceptibility in the presence of antioxidants was seen in *Candida krusei* cells exposed to the polyunsaturated fatty acid, γ-linolenic acid (Jamiu *et al*., 2021).

**Figure 2.**
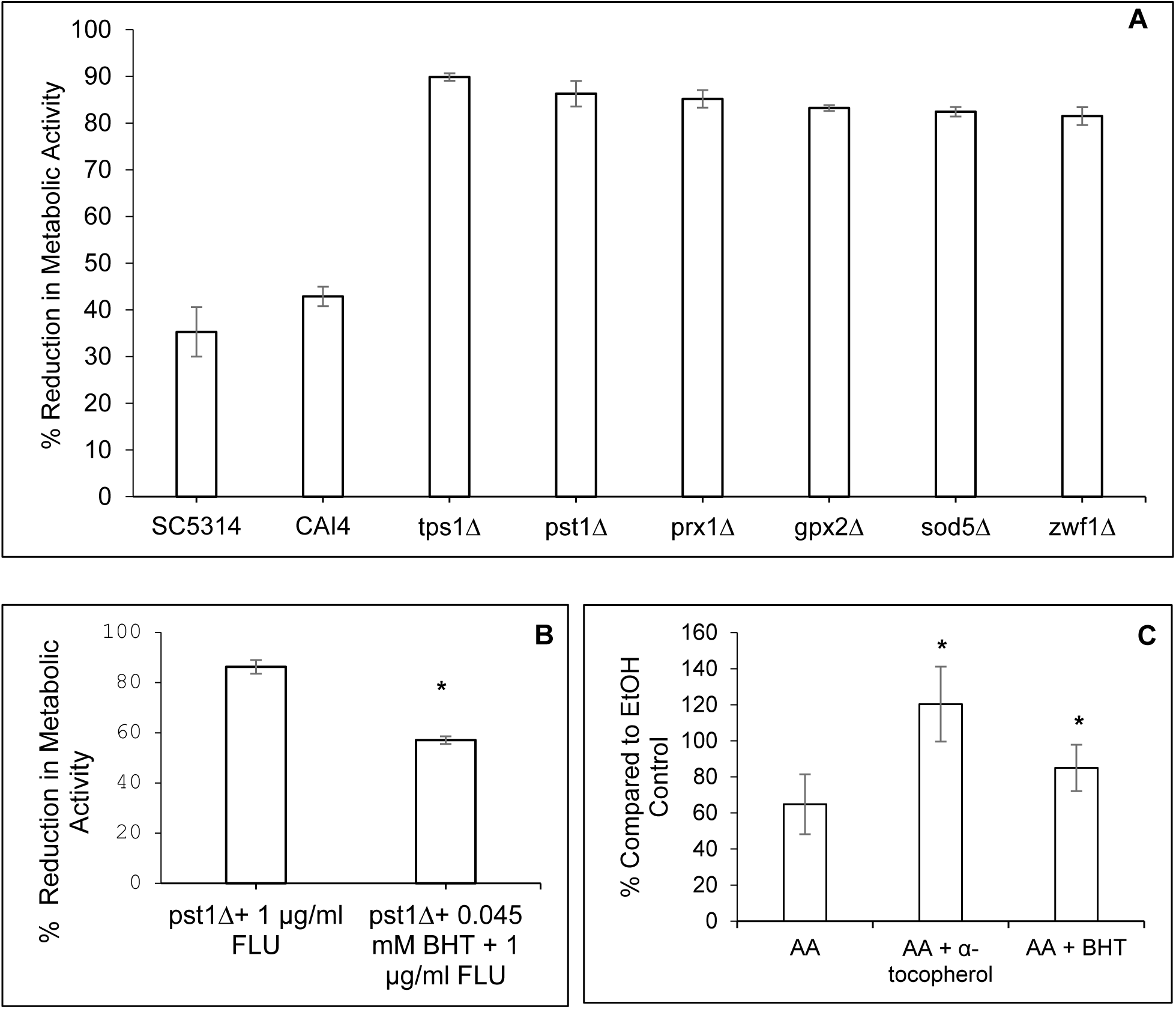
Role of arachidonic acid (AA) in response to fluconazole-induced oxidative stress. A) Effect of fluconazole on biofilm formation of selected heterozygous mutants. B) Effect of fluconazole on *PST1* heterozygous mutants (*pst1Δ/PST1*) and the reversals in the presence of butylated hydroxytoluene (BHT). C) Biomass of *Candida albicans cdr1*Δ/Δ in the presence of 0.1 mM arachidonic acid (AA) and antioxidants. Percentage biofilm biomass after 6 h of *cdr1*Δ/Δ in the presence of 0.1 mM arachidonic acid compared to ethanol control (0.076%). In addition, 0.5 mM α-tocopherol and 0.045 mM butylated hydroxytoluene (BHT) were added, respectively. Values are the mean of three independent experiments, and the standard deviation is indicated by the error bars.*Significantly different from cdr1Δ/Δ with only arachidonic acid (\**P* < 0.05).

Since Cdr1p is a promiscuous transporter with proclivity for hydrophobic compounds (Prasad *et al*., 2015; Prasad *et al*., 2019), we also investigated the role of Cdr1p in protecting *C. albicans* from arachidonic acid-induced oxidative stress. Addition of arachidonic acid inhibited biofilm formation by *cdr1*Δ/Δ (Figure 2c), suggesting that the absence of *CDR1* may abrogate the ability of cells to export arachidonic acid-containing lipids or their oxidation products. To test this, *cdr1*Δ/Δ was exposed to arachidonic acid as well as BHT and α-tocopherol. Due to the reducing capabilities of α-tocopherol which interferes with tetrazolium salt assays such as the XTT assay (Lim *et al*., 2015), the effect of supplementation with antioxidants on biofilm formation was only measured by crystal violet assay. The addition of both antioxidants reduced the inhibitory effect of arachidonic acid on *cdr1*Δ/Δ biofilm formation, indicating that the reduction in biomass of *cdr1*Δ/Δ may be due to oxidative stress. The effect of α-tocopherol was more pronounced and since α-tocopherol localises to membranes (as opposed to the more general antioxidant properties of BHT), it may indicate that the oxidative stress experienced by *cdr1*Δ/Δ is mainly due to lipid peroxidation in the membranes (Li *et al*., 2015; Yehye *et al*., 2015; Yin *et al*., 2011). These results indicate that the systems that protect *C. albicans* from fluconazole-induced oxidative stress are suppressed by arachidonic acid and that this may be a contributing mechanism to the observed increased fluconazole susceptibility.

### Arachidonic acid increases efflux pump expression, but reduces activity

According to data by Kuloyo and co-workers (2020) fluconazole increased the expression of *MDR1*, *CDR1,* and *CDR2* (Table 4). Similar results were also observed by Shi and co-workers (2019) upon the addition of fluconazole to *C. albicans* biofilms, indicating the role of these drug efflux pumps in the response to fluconazole. Interestingly, arachidonic acid induced the further upregulation of *CDR1* and *CDR2* as well as genes encoding other ABC transporters and the major-facilitator-superfamily transporter, *FLU1*.

**Table 4.**
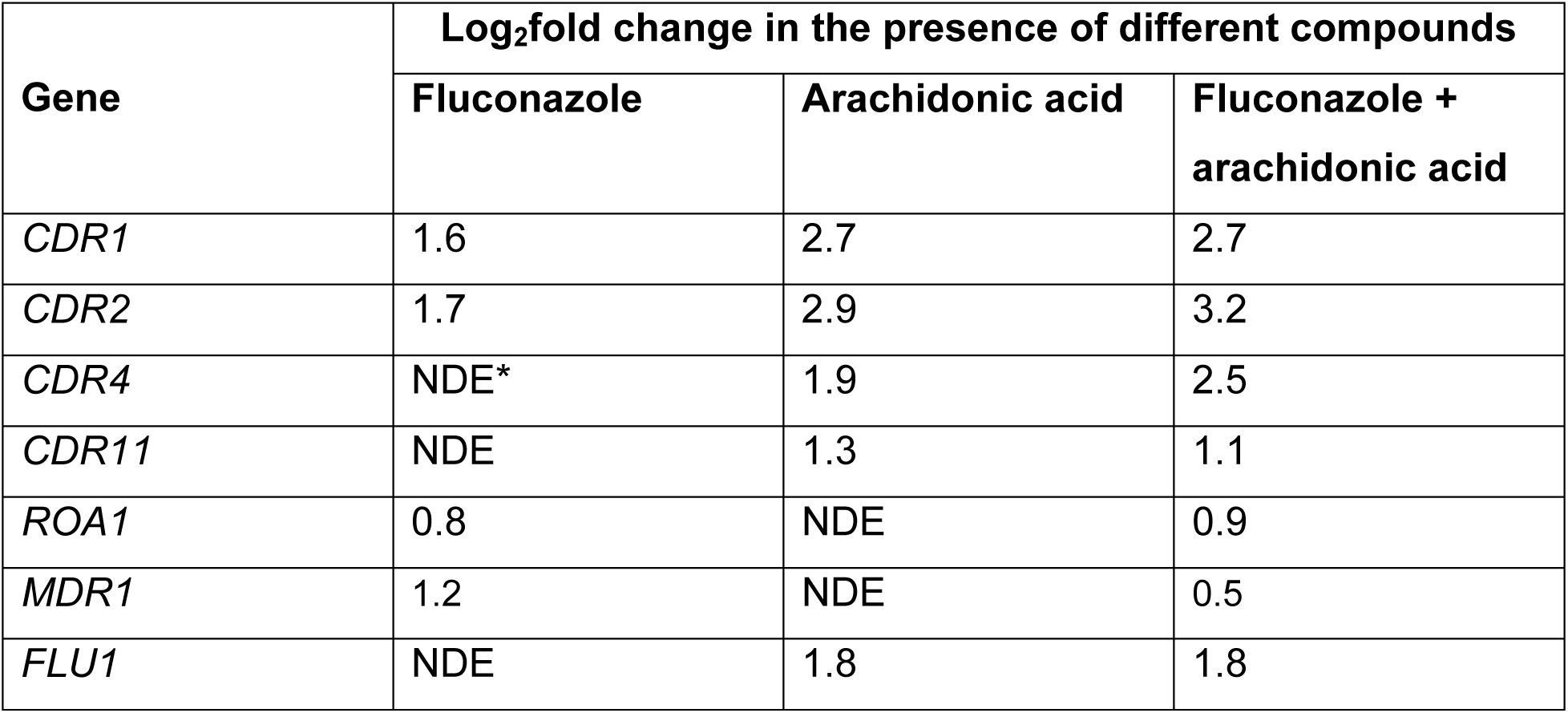
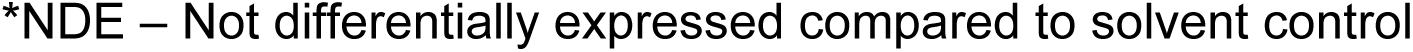
Differential expression of efflux pump genes in *Candida albicans* exposed to fluconazole, arachidonic acid, or a combination of the two compounds.

| Gene | Log<sub>2</sub>fold change in the presence of different compounds |  |  |
| --- | --- | --- | --- |
|  | Fluconazole | Arachidonic acid | Fluconazole + arachidonic acid |
| *CDR1* | 1.6 | 2.7 | 2.7 |
| *CDR2* | 1.7 | 2.9 | 3.2 |
| *CDR4* | NDE* | 1.9 | 2.5 |
| *CDR11* | NDE | 1.3 | 1.1 |
| *ROA1* | 0.8 | NDE | 0.9 |
| *MDR1* | 1.2 | NDE | 0.5 |
| *FLU1* | NDE | 1.8 | 1.8 |

\*NDE – Not differentially expressed compared to solvent control

Considering the reported increase in fluconazole susceptibility in the presence of arachidonic acid (Ells *et al*., 2009), the increase in expression of efflux pump-genes was surprising. However, since the observed increase in expression may not reflect an increase in protein levels, the level of Cdr1p was evaluated using a strain with GFP fused to *CDR1* (AsCa1). As can be seen in Figure 3, low fluorescence was observed in control cells, but increased fluorescence was observed in biofilms grown in the presence of 1 mM arachidonic acid, indicating an increase in Cdr1p. Interestingly, the fluorescence in the presence of arachidonic acid was not localised on the cell periphery as expected for active efflux pumps, but in the cytoplasm, suggesting delocalisation of Cdr1p.

**Figure 3.**
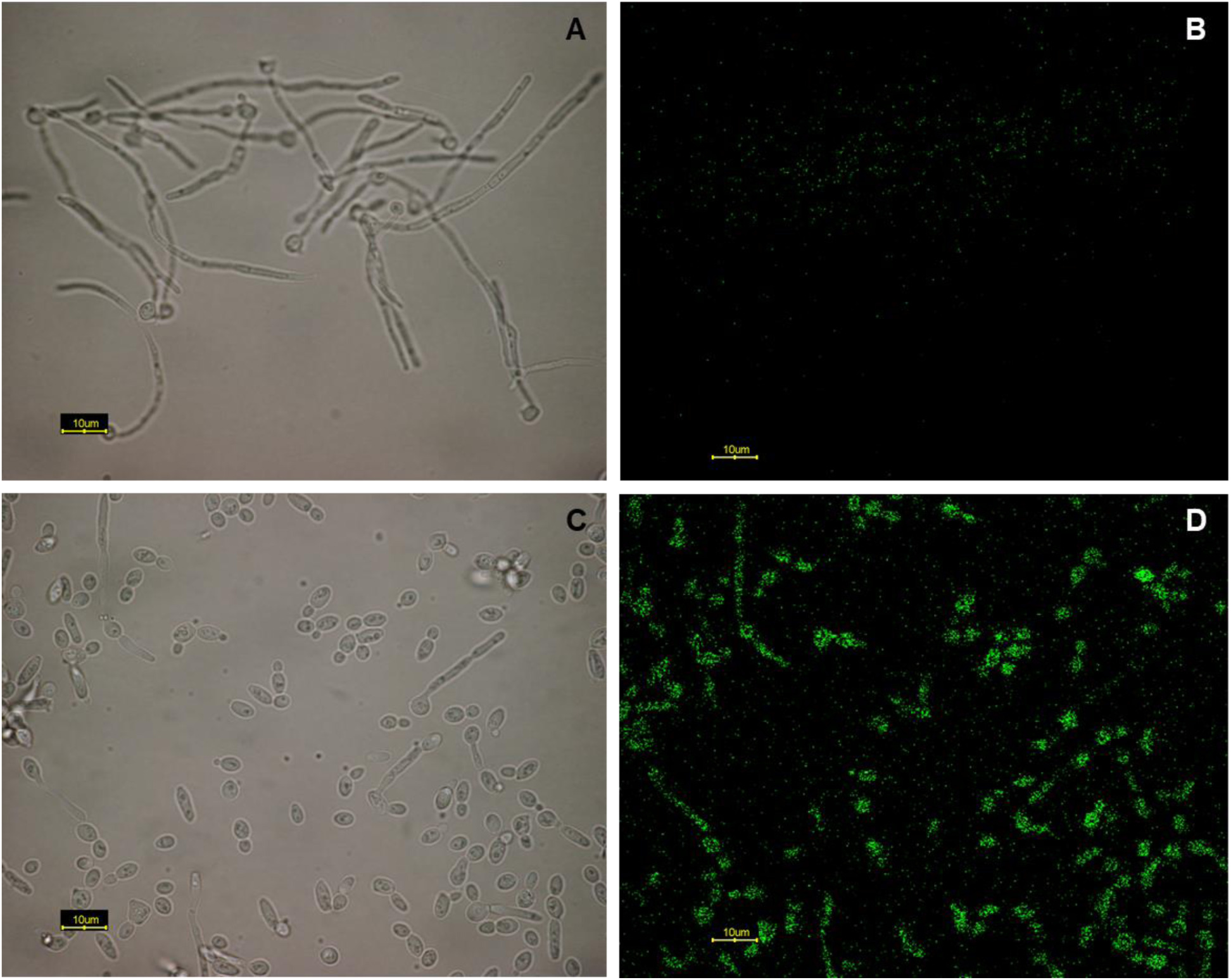
Fluorescence of *Candida albicans* CDR1-GFP (strain AsCa1) with exposure to ethanol control and 1 mM arachidonic acid. **A** and **C** represent light micrographs, whereas **B** and **D** are corresponding fluorescent micrographs. **A** and **B** – AsCa1 exposed to ethanol; **C** and **D** – AsCa1 exposed to 1 mM arachidonic acid. Scale bars represent 10 μm.

Since it is possible that delocalisation of Cdr1p may occur, which will influence efflux activity, we determined the activity of Cdr1p. To evaluate this, the efflux of R6G was followed in the presence of arachidonic acid in two ways. Firstly, 6-h biofilms were allowed to form in the presence of arachidonic acid (added at time zero) and secondly, biofilms were allowed to form for 6 h in the absence of arachidonic acid and then exposed to arachidonic acid for 1 h prior to measurement of R6G-efflux.This would indicate if arachidonic acid is required during biofilm formation to elicit an effect on Cdr1p, or if short-term treatment could elicit this effect. As can be seen in Figure 4, the addition 1 mM arachidonic acid at either time point resulted in significant inhibition of drug efflux activity. Similar results regarding inhibition of efflux activity by γ-linolenic acid was observed in *C. krusei* (Jamiu *et al.,* 2021).

**Figure 4.**
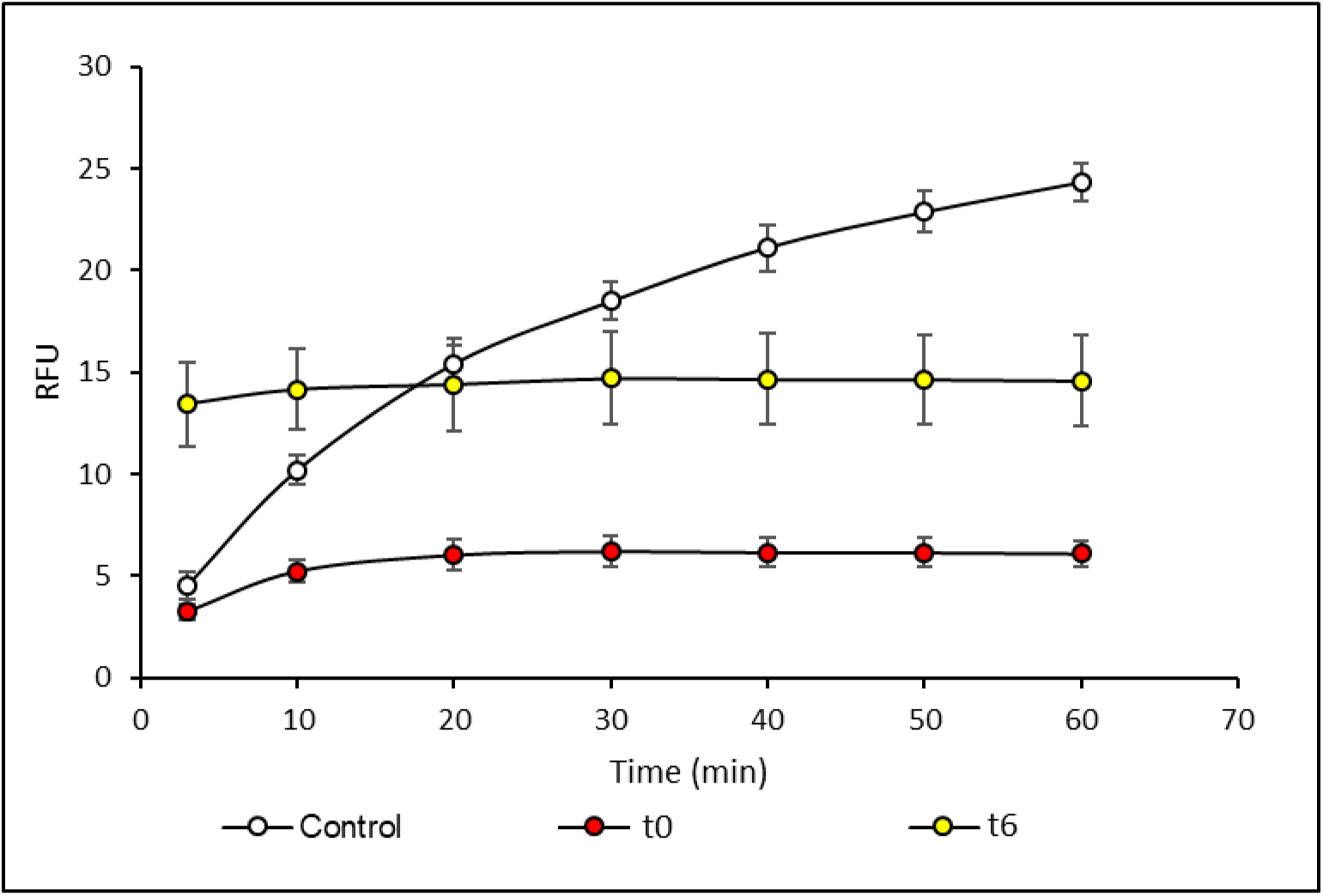
Rhodamine 6G-efflux of *Candida albicans* in the presence of ethanol or arachidonic acid. Y-axis represents relative fluorescent units (excitation/emission: 530/590 nm). Control – biofilms exposed to solvent (ethanol); t0 – 1 mM arachidonic acid added prior to biofilm formation; t6 – 1 mM arachidonic acid added after biofilm formation (6 h) for 1 h .

### Arachidonic acid causes changes in phosphorylation of Cdr1p

Cdr1p is subject to post-translational modification through phosphorylation of the N-terminal extension (Tsao *et al*., 2016). Thus, we investigated the effect of arachidonic acid and fluconazole on phosphorylation of proteins.

Differential production of Cdr1p phosphopeptides was observed (Table 5) in cells exposed to either fluconazole or arachidonic acid, especially within the N-terminal extension. Unexpectedly, in the presence of fluconazole, less peptides containing these phosphorylated amino acids were detected compared to the control cells, while the addition of arachidonic acid increased the level of phosphorylation compared to the control cells at S7 and S12 and did not cause a decrease in phosphorylation at the other amino acids.

**Table 5.** Differential phosphorylation of amino acids of Cdr1p in response to fluconazole or arachidonic acid.

| Phosphosite | Log<sub>2</sub>fold change* in phosphopeptide abundance |  |
| --- | --- | --- |
|  | Fluconazole | Arachidonic acid |
| S7 | NDP** | 2.0 |
| S8 | -0.7 | NDP |
| S12 | NDP | 1.1 |
| S54 | -2.2 | NDP |
| T51 | -1.6 | NDP |

\*P values ≤0.05
\*\* NDP – Not differentially phosphorylated in comparison to the control

The exact effect of changes in phosphorylation of Cdr1p is unknown but multiple studies have highlighted the effects changes in phosphorylation may have on the overall fold of a protein. These effects include the disruption or creation of secondary structural elements such as α-helixes and β-pleated sheets, the folding or unfolding of disordered regions, changes in the *cis-trans* orientation of amino acid side chains and the association or dissociation of multiple peptide assemblies. These changes in return can cause large scale shifts in the overall protein fold, halting, stopping or changing the activity of the protein (Gaffarogullari *et al*., 2011; Kobashigawa *et al*., 2015; Newcombe *et al*., 2022). Crucially, it is known that specific combinations of phosphorylation events can have distinct and diverse consequences for the activity of a protein (Bilbrough *et al*., 2022). Thus, although the impact of the specific phosphorylation events on Cdr1p structure and function is currently unknown, it may be speculated that these changes could impact the activity of the efflux pump, contributing to increased fluconazole susceptibility in the presence of arachidonic acid.

## Conclusions

This study provides evidence that the host-derived polyunsaturated fatty acid, arachidonic acid, can modulate multiple fluconazole resistance mechanisms, including upregulation of *ERG11* and oxidative stress responses in *C. albicans* biofilms, contributing to increased susceptibility. Notably, although arachidonic acid increases *CDR1* transcription and Cdr1p expression, it paradoxically reduces efflux activity, likely through mislocalisation of the Cdr1p, and altered phosphorylation patterns. This indicates that expression levels of *CDR1* may not always correlate to fluconazole susceptibility. Similar findings were also observed for other polyunsaturated fatty acids and in other pathogenic yeasts (Thibane *et al*., 2012; Jamiu *et al*., 2021) and highlight the important and diverse influences of polyunsaturated fatty acids in the response to fluconazole. This work opens avenues for further exploration of lipid-mediated modulation of antifungal responses.

